# TomoNet: A streamlined cryoET software pipeline with automatic particle picking on flexible lattices

**DOI:** 10.1101/2024.02.17.580557

**Authors:** Hui Wang, Shiqing Liao, Xinye Yu, Jiayan Zhang, Z. Hong Zhou

## Abstract

Cryogenic electron tomography (cryoET) is capable of determining *in situ* biological structures of molecular complexes at near atomic resolution by averaging half a million subtomograms. While abundant complexes/particles are often clustered in arrays, precisely locating and seamlessly averaging such particles across many tomograms present major challenges. Here, we developed TomoNet, a software package with a modern graphical user interface to carry out the entire pipeline of cryoET and subtomogram averaging to achieve high resolution. TomoNet features built-in automatic particle picking and 3D classification functions and integrates commonly used packages to streamline high-resolution subtomogram averaging for structures in one-, two- or three-dimensional arrays. Automatic particle picking is accomplished in two complementary ways: one based on template matching and the other employing deep learning. TomoNet’s hierarchical file organization and visual display facilitate efficient data management as required for large cryoET datasets. Applications of TomoNet to three types of datasets demonstrate its capability of efficient and accurate particle picking on flexible and imperfect lattices to obtain high-resolution 3D biological structures: virus-like particles, bacterial surface layers within cellular lamellae, and membranes decorated with nuclear egress protein complexes. These results demonstrate TomoNet’s potential for broad applications to various cryoET projects targeting high-resolution in situ structures.

## INTRODUCTION

Single-particle cryogenic electron microscopy (cryoEM) is employed to elucidate atomic-level structures of purified biological complexes. This methodology adheres to a standardized and well-established workflow supported by advanced software packages such as Relion^1^ and cryoSparc^2^. In parallel, cryogenic electron tomography (cryoET), coupled with subtomogram averaging (STA), expands the investigative scope to encompass heterogeneous macromolecules in their native context^3–10^. To enhance the resolution of subunits within *in situ* macromolecules, subtomograms (*i.e.,* particles) are extracted from each tomogram and then subjected to 3D alignment and averaging, thereby improving signal-to-noise ratio. Notably, STA has achieved resolutions up to sub-3 Å for *in situ* structures of large cellular complexes such as ribosomes, approaching the capabilities of single-particle cryoEM methodologies^11–14^.

The workflow for cryoET and STA typically involves five key components across specific software packages. In cryoET preprocessing, dose fractionated frames are collected from an electron microscope, undergo motion correction, organized, and then assembled into individual tilt series. In tomogram reconstruction, three-dimensional reconstructions are generated from those tilt series. In particle picking, particles of interest are identified and extracted from tomograms. Complexity varies based on the diverse and intricate nature of *in situ* cellular samples and their unique configurations. Many packages include their own particle picking methods, such as oversampling using a supporting geometry in Dynamo^15,16^, template matching in emClarity^17^ and machine learning in crYOLO^18^. In 3D refinement and classification, particles are iteratively classified and refined to obtain a final structure at sub-nanometer or near atomic resolution, which has been demonstrated by software packages like Relion^13,19^, emClarity^17^, EMAN2^4^ and Warp^20^. Finally, activities in post-processing include map sharpening, Fourier shell correlation (FSC) calculation, visualization by placing averaged maps back into the original tomogram, etc. Users often need to navigate between several specialized software packages for optimal results, which often demands a certain level of computational proficiency that poses a barrier for many.

The method for particle picking varies on a case-by-case basis, dictated by the characteristics of *in situ* cellular samples. In the early works of STA, manual particle picking was employed, particularly when aiming for resolutions between 20-50 Å with a maximum of several hundred particles^21–23^. However, for biological samples exhibiting periodic structures, oversampling on specified geometry was leveraged to significantly reduce the labor associated with acquiring enough particles for improved resolutions. For instance, HIV virus-like particles (VLPs) adopt a hexagonal Gag protein lattice in its sphere-like configuration^16^. Other examples include the Marburg Virus^24^, Herpes simplex virus^25^, and the Coat protein complex II^26^, all of which contain lattice-like arrangements with repeating subunits that could benefit from particle picking automation when performing cryoET data processing. With an increasing demand for automation to enhance efficiency with minimal manual intervention, template matching has emerged as a popular method for automatic particle picking, relying on a user-provided reference map^17,27^. Simultaneously, convolutional neural networks have shown promising results for cryoET automatic particle picking given its capacity to analyze three-dimensional feature maps and autonomously identify prominent features within specific samples^28–31^. These machine learning approach typically operate template-free and often obviates the need for human annotation^32^.

The expanding array of specialized software tools designed for specific tasks posts a critical need for seamless software integration within the cryoET workflow. Transitioning between various software packages can be a cumbersome process. Remarkably, recent initiatives have made notable progress in tackling this integration challenge. For example, TomoBEAR^33^ offers an integrated solution, while ScipionTomo^34^ and nextPYP^35^ provide a comprehensive web-based platform for managing various tasks in the cryoET pipeline. Notably, none of these packages takes specific advantage of the fact that abundant complexes exist in arrays of some sort, albeit with imperfections, variability, or flexibility.

In this context, we have developed TomoNet, a software package designed for streamlining the cryoET and STA data processing workflow, with a modern GUI (Figure 1 and Figure 2). Our methodology employs a geometric template matching approach, rooted in the concept of “Auto Expansion”, which serves as a general particle picking solution for biological complexes organized in flexible, variable, or imperfect arrays. TomoNet is also powered by a deep learning-based solution to automate particle picking, which only needs 1-3 tomograms with known particle locations as ground truth for model training. Importantly, while TomoNet is particularly powerful for locating and averaging particles arranged on flexible or imperfect lattices, it can be applied to a broader range of particle types, offering a more generalizable trained model. These methods significantly diminish the need for manual inputs, and their outcomes can be seamlessly imported into Relion for subsequent high-resolution 3D classifications and refinements. We demonstrate the capabilities of TomoNet by applying it to three datasets with distinct protein lattice types, highlighting its accuracy and efficiency in identifying particles across diverse scenarios.

**Figure 1.**
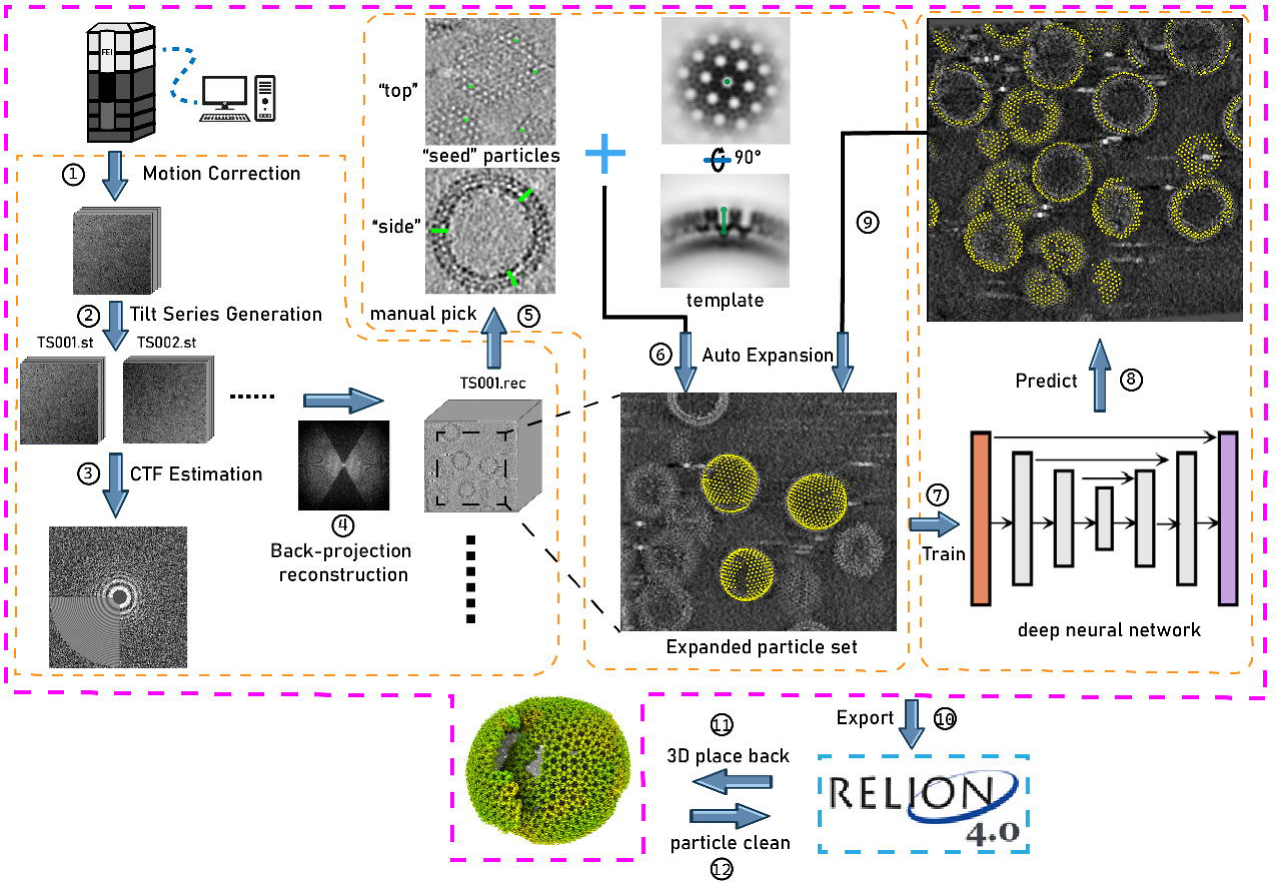
Illustration of TomoNet’s comprehensive pipeline for cryoET and STA. The pink border encloses the sequential functions implemented in TomoNet, and they can be subdivided into three principal segments, delineated by the orange borders. These segments include tomogram preparation on the left, template matching-based particle picking “Auto Expansion” in the center, and deep learning-based automatic particle picking on the right.

**Figure 2.**
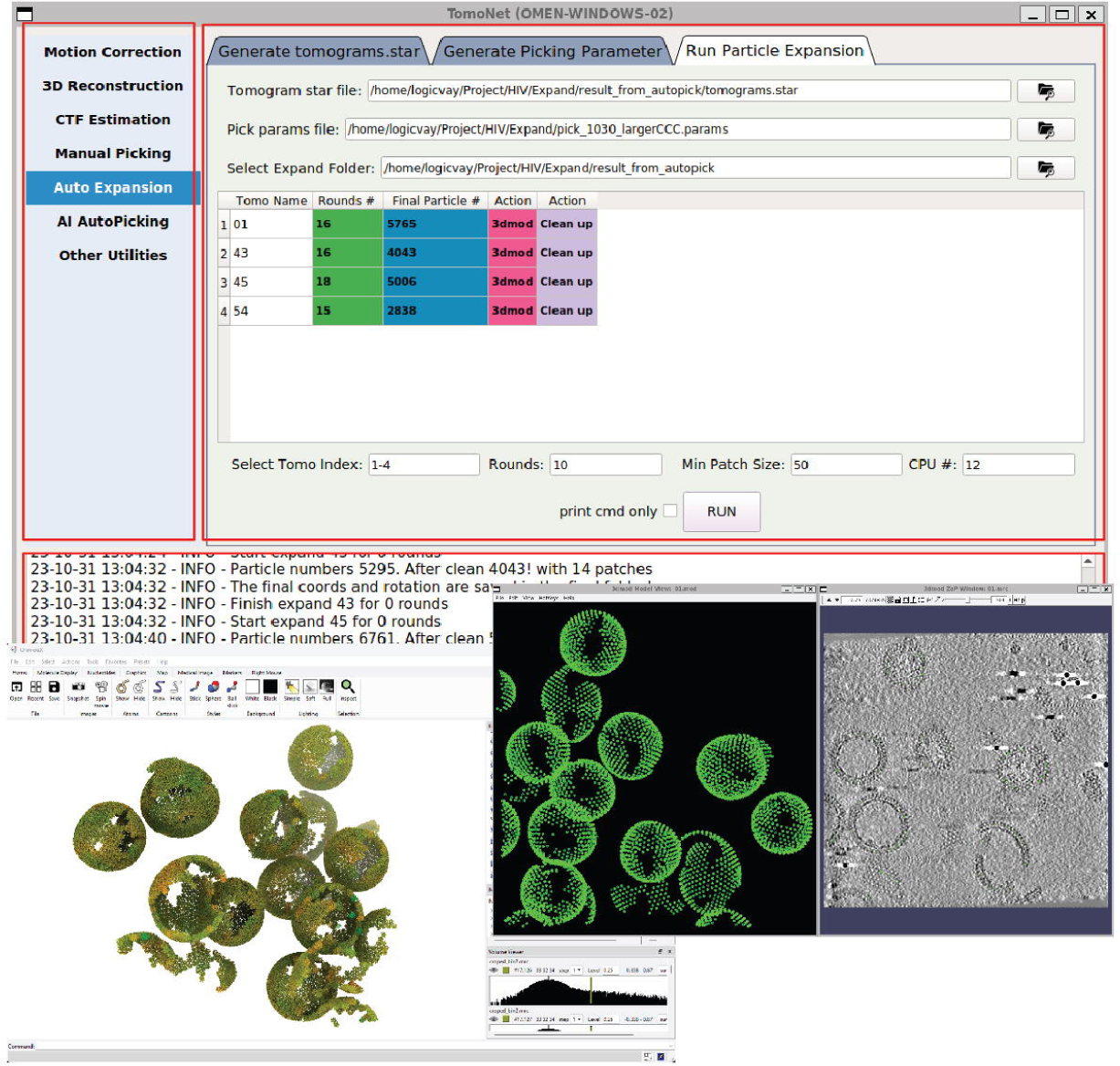
A screenshot of TomoNet GUI. The TomoNet GUI contains three main areas: the menu bar (top left), the input and operate area (top right), and the log window (bottom). Bottom left: results generated by the “3D Subtomogram Place Back” function can be visualized in ChimeraX. Bottom right: intermediate results of picked particles viewed with IMOD.

## RESULTS

### Overall design of TomoNet

TomoNet is a Python-based software package that integrates commonly used cryoET packages to streamline the cryoET and STA pipeline, with a particular emphasis on automating particle picking of lattice-configured structures and cryoET project management. As shown in the main menu and the entire TomoNet pipeline (Figure 1 and Figure 2), after data collection from electron microscopy, TomoNet can perform motion correction with integration of MotionCorr2^36^; tilt series assembly and tomogram reconstruction with integration of IMOD^37^ and AreTomo^38^; CTF estimation with integration of CTFFIND4^39^; manual particle picking with IMOD; particle picking using built-in geometric template matching-based algorithms with integration of PEET^40^; automatic particle picking using built-in deep learning-based algorithms; 3D classification/particle cleaning and subtomograms placing back with built-in algorithms. This design also allows on-the-fly tomogram reconstruction processing during data collection, which facilitates a quick quality check. TomoNet generates particle picking results in STAR format^41^, which can be incorporated into Relion for high-resolution 3D refinement. It can also read Relion results in STAR format for particle cleaning and subtomograms placing back (Figure 1).

### Particle picking with “Auto Expansion”

The “Auto Expansion” module is based on template matching and uses cross-correlation coefficient as a selection criterion, with a design to pick particles on flexible lattices with minimal manual inputs, its basic concept is elucidated in Figure 3. These particles exist in array-like configurations and manifest as flexible, partial, and imperfect lattices in one, two and three dimensions (1-3D). Examples are abound: microtubule doublets, ubiquitous in most cells, consist of 96 nm axonemal 1D translational repeat units^22,42^ (1D rotational lattice); HIV VLPs^43^ and surface layer (S-layer) lattice of prokaryotic cells^44,45^ are composed of hexametric subunits (2D lattice); paraflagellar rod of protozoan species is organized into para-crystalline arrays in its distal zone^21^ (3D lattice). In TomoNet, each of these isolated lattice densities is called a patch, within which all subunits of the complex are connected. For instance, Figure 3 illustrates two patches with different sizes.

**Figure 3.**
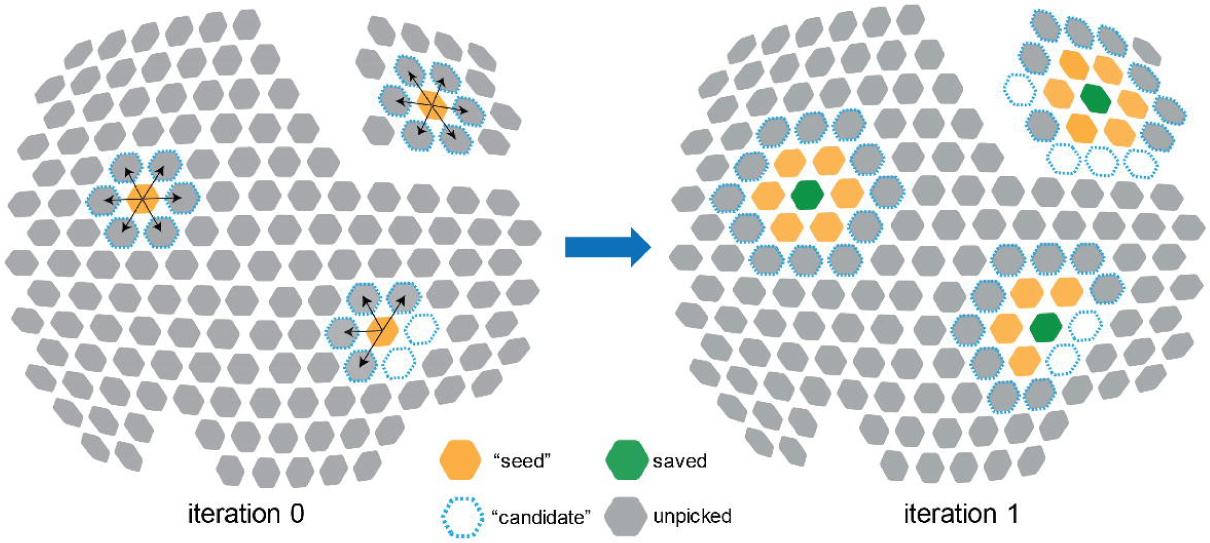
Illustration of the first two iterations of “Auto Expansion“ particle picking. There are two patches of a hexagonal lattice with individual particles represented by solid hexagons. At iteration 0, 18 “candidate” particles (dashed blue) were selected from the neighbors of 3 “seed” particles (orange). 14 good particles remained and will serve as “seed” particles in iteration 1, and 3 “seed” particles in iteration 0 were saved in the final particle set (green). At iteration 1, 35 “candidate” particles were selected from the neighbors of 14 “seed” particles. 29 good particles remained and will serve as “seed” particles in iteration 2, and 14 “seed” particles were saved in the final particle set. “Auto Expansion” is an iterative process and will stop when no “candidate” can be detected.

“Auto Expansion” is an iterative process; each iteration expands the particle set by adding more unpicked ones. To initiate “Auto Expansion”, users need to prepare a few “seed” particles that sparsely distribute across all observed patches. Typically, the numbers of such “seed” particles per tomogram range from 20 to 200, which depends on the number and size of patches in the input tomogram. Then, “Auto Expansion” iteratively expands the “seed” particle set to a final particle set that contains all particles on given flexible lattices, following three steps for each iteration (Figure 3). Firstly, potential particles adjacent to each “seed” particle are calculated and selected as “candidate” particles. Secondly, these “candidate” particles undergo alignments to a user-provided reference and are evaluated based on cross-correlation coefficient, such that “wrong” particles with low cross-correlations are excluded. Thirdly, qualified “candidate” particles are added to the particle set and become “seed” particles for the next iteration. During this process, only unpicked ones can be considered as “candidate” particles, and “Auto Expansion” stops either when no “candidate” particles are detected or when the user-defined maximum iteration number is reached. Doing this allows for an exhaustive exploration of particles on given lattices following their assembly topology with no restriction on geometry and outputs a final particle picking result (Figure 2).

Compared with conventional template matching methods, “Auto Expansion” incorporates prior knowledge of lattice configuration to iteratively guide the search for “candidate” particles, *i.e.*, unpicked particles following user-defined paths, as detailed in the Method section and TomoNet’s user manual. Thus, “Auto Expansion” significantly reduces computational complexity by searching in the regions of interest only, with restricted angular and translational search ranges defined by users. As a result, it reduces the number of incorrectly picked particles. Notably, “Auto Expansion” potentially works for any flexible, imperfect, or variable lattices in 1D, 2D and 3D and has no intrinsic size limit of subunits.

### Automatic particle picking by deep learning

The “AI AutoPicking” module is designed for automatic particle picking using supervised machine learning, which employs a U-net convolutional neural network for model training. There are three main steps in “AI AutoPicking”: training data preparation, neural network training, and particle coordinate prediction, as detailed in the Method section (Figure 4). It only requires an input training dataset consisting of 1-3 tomograms paired with their corresponding particles coordinate files. The trained model can then be applied on the entire tomography dataset and output predicted particles for each tomogram.

**Figure 4.**
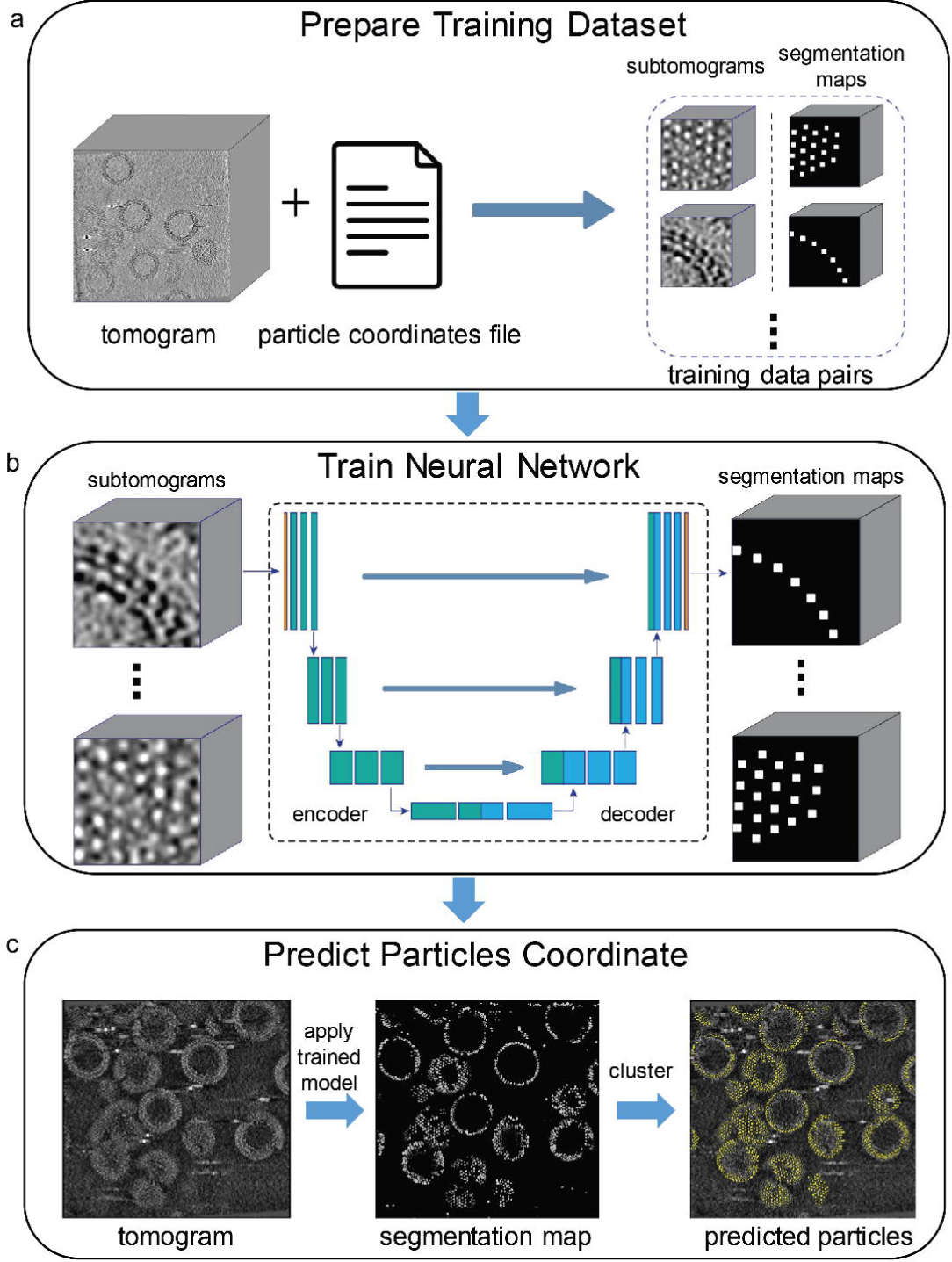
Illustration of “AI AutoPicking“ process consisting of three steps. The HIV dataset was used for this illustration, and the particles refer to Gag hexamers. **a,** Training dataset preparation. Using the user-provided tomograms with associated particle coordinate files, subtomograms containing particle densities were extracted. For each subtomogram, TomoNet generated a segmentation map based on the coordinates of particles, where the voxels near a particle’s center are shown as white and the others as black. **b,** Neural network training. The generated subtomograms and segmentation maps were used as the input and output to train the convolutional neural network in learning how to segment out particle densities. **c,** Particle coordinate prediction. Firstly, TomoNet applied the trained neural network model to unseen tomograms and generated associated predicted segmentation maps. Then, the particle coordinate information was obtained from the segmentation maps using clustering algorithms.

Essentially, the neural network in “AI AutoPicking” is trained as a voxel-wise binary classifier, which determines whether a voxel in density maps is part of a particle (Figure 4b). To prepare for training, data pairs (ground truth) consist of extracted subtomograms coupled with their associated segmentation maps, within where each particle is labeled by a cube near its center (Figure 4a). The trained neural network model can be applied on other tomograms to perform particle segmentation. Finally, the particles coordinate information can be retrieved from the predicted segmentation maps (Figure 4c).

### 3D classification using TomoNet

In addition to the above two commentary modules for particle picking, TomoNet allows users to eliminate “bad” particles based on user-defined geometric constraints, which could serve as 3D classification during high-resolution particle refinements. Lattice variation in cryoET data has multiple plausible causes. Biologically, particles may be incomplete near the lattice edge due to paused biology assembly process^46^. Experimentally, lattices tend to become flattened near the air-water interface of the sample during imaging. These variabilities pose challenges for 3D classification in the process of high-resolution STA, making it difficult to exclude “bad” particles that exhibit unexpected coordinates and orientations assignment as subunits of lattices (Supplementary Movie 1).

Removing these “bad” particles is necessary for achieving better resolutions^47^. To accomplish this, TomoNet assesses each particle by counting its neighboring particles and calculating the averaged tilt angle to these neighbors to represent local surface curvature of a lattice. TomoNet identifies particles with too few neighbors or large tilt angles to their neighbors as “bad” particles since they potentially deviate from the lattice configuration. This step can be integrated into high-resolution refinement in Relion, providing an alternative 3D classification method based on analyzing spatial relationships between particles.

### Application to *in situ* viral protein arrays: the matrix protein lattice in HIV VLPs

To validate TomoNet as an integrated high-resolution cryoET and STA pipeline and an efficient particle picking tool, four tomograms were processed from the HIV-1 Gag dataset which resolved the Gag hexamer structure at 3.2 Å resolution. Motion corrected images underwent tilt series assembly, CTF estimation, and tomographic reconstruction using TomoNet. Within these tomograms, the VLP hexagonal lattice and its building blocks were observed, and some of these observed VLPs exhibited sphere-like geometry (Figure 5a).

**Figure 5.**
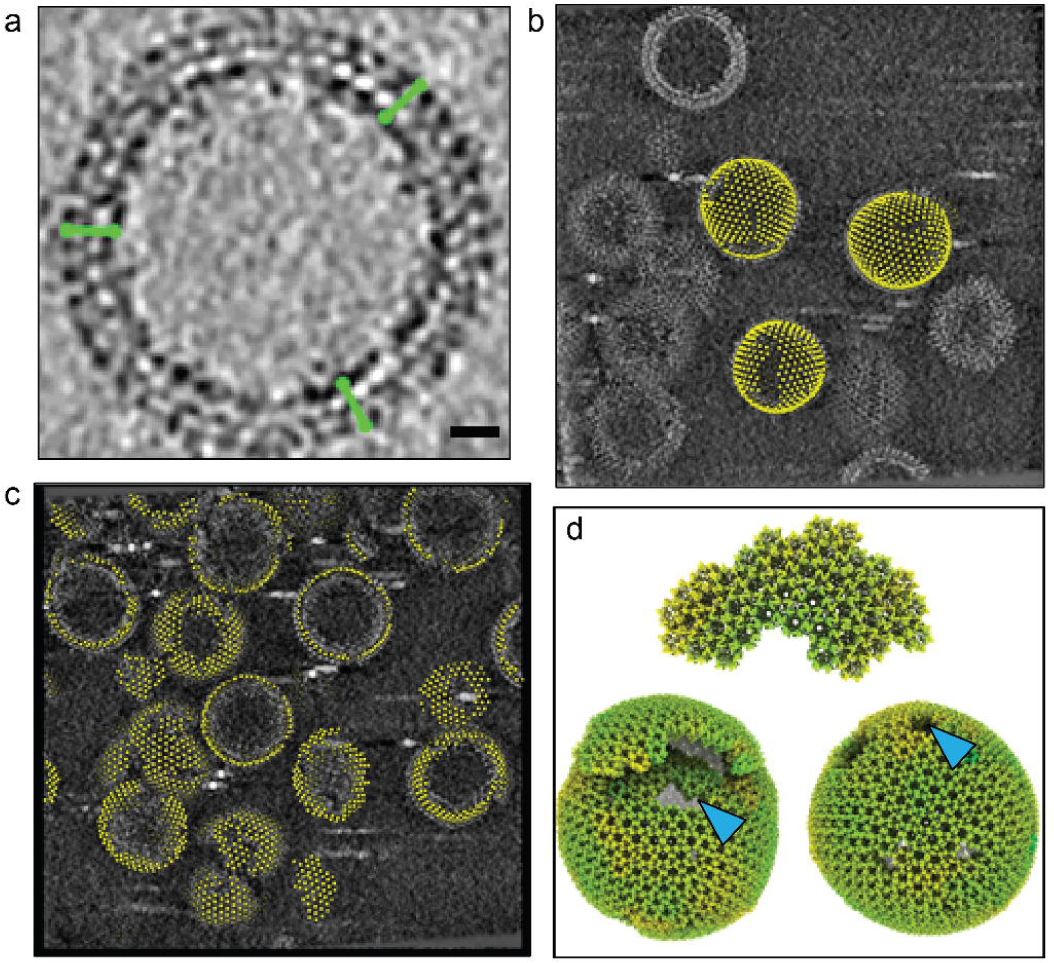
TomoNet application to arrays of matrix protein in HIV VLPs. **a,** Illustration of picked “seed” particles on a spherical VLP. Green segments represent the particles’ Y-axis. Scale bar is 20 nm. **b,** “Auto Expansion” result on three VLPs within tomogram TS_01, with yellow dots representing the center of the hexamer subunits. **c,** “AI AutoPicking” particle prediction result of tomogram TS_45 shows its ability to pick particles on all lattices of different sizes and shapes. **d,** Visualization of three different variations of the HIV Gag lattices generated by placing back averaged structures, two exhibiting a spherical shape, and one presented as a fragment. Blue arrows indicate defects in the lattice.

As detailed in the Method section, a combination of “Auto Expansion” and “AI AutoPicking” was applied on the above four tomograms; as a result, particles were readily picked on all the observed lattice patches (Figure 5b, c). Then, these picked particles were imported to Relion to perform high-resolution particle refinements, resulting in a final reconstruction of the Gag hexamer structure (Figure 6).

**Figure 6.**
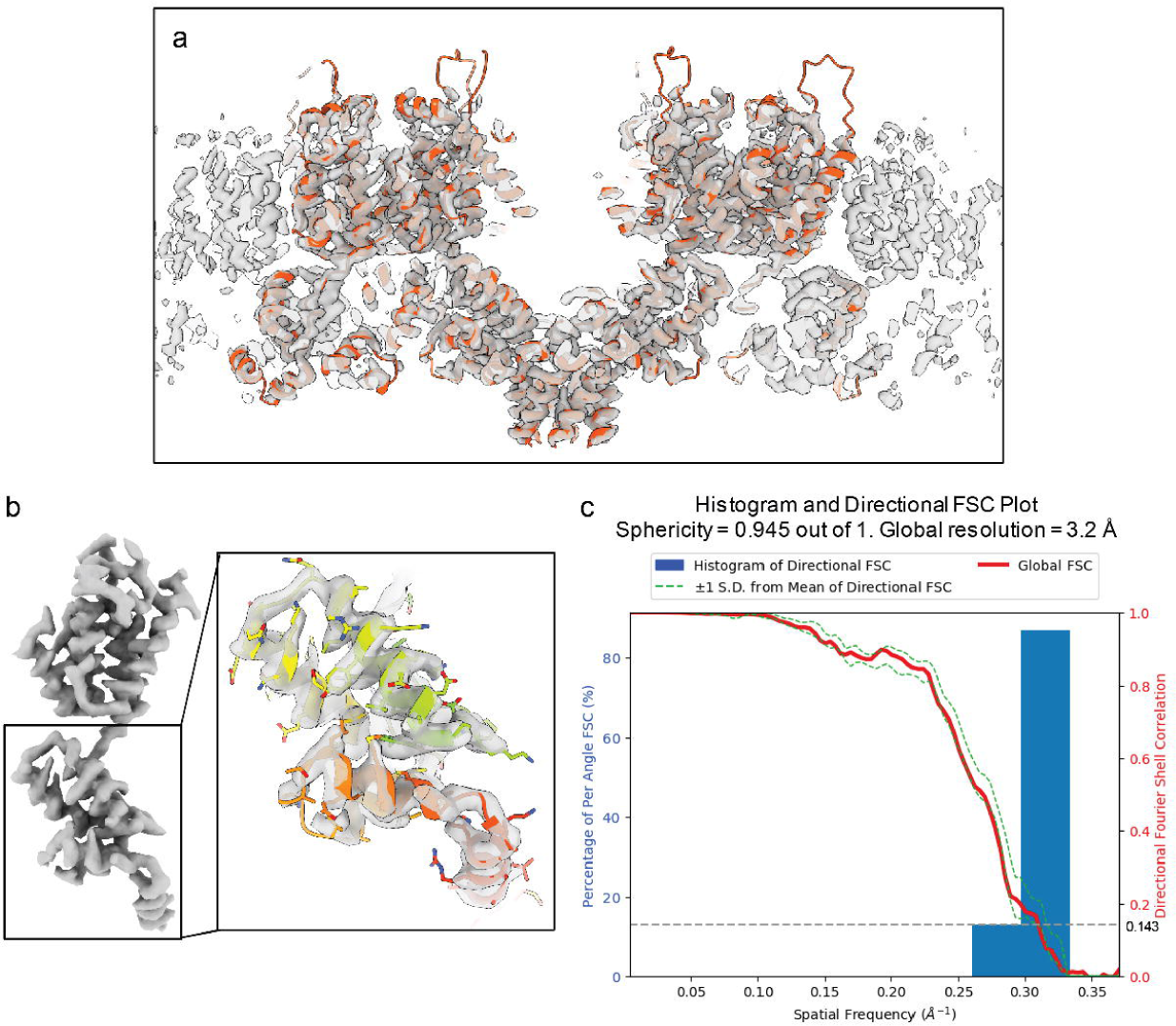
Final map resolution of HIV Gag hexamer. **a,** Final reconstruction of Gag hexamer (grey) fitted with the atomic model (PDB: 5l93). **b,** One segmented Gag monomer structure, inset shows a closer view of carboxy-terminal domain overlay with the atomic model. **c,** Directional Fourier shell correlation (FSC) curves for the STA of Gag hexamer structure, with a global resolution at 3.2 Å.

Using the “3D subtomogram place back” function in TomoNet, 3D visualizations were generated to illustrate the *in situ* assembly of the VLP lattices (Figure 5d and Figure 7). All VLP lattices with various sizes and shapes were captured even with irregular shapes (Figure 7e and Supplementary Movie 2), demonstrated TomoNet’s particle picking ability on flexible lattices. Lattice defects on each VLP were also identified consistent with previous studies^48^, enhancing the understanding of lattice assembly mechanisms^49^.

**Figure 7.**
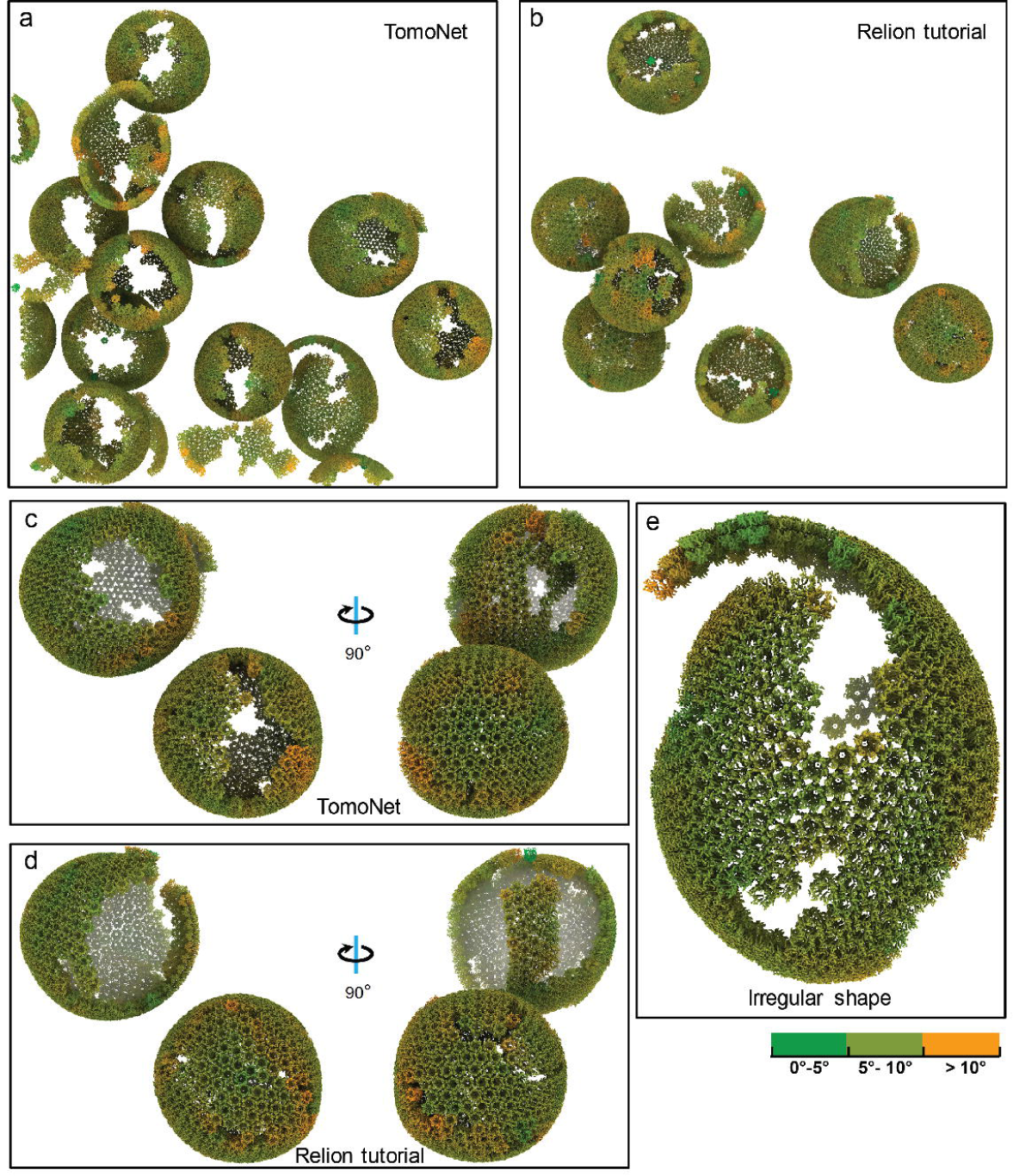
Comparative visualization of lattices obtained from TomoNet and Relion tutorial. **a, b,** Visualized comparison of particles used in TomoNet and Relion tutorial within tomogram TS_01. TomoNet can pick particles not only on a sphere-like lattice but also on others with random shapes. **c, d,** A comparison of particle picking results on two sphere-like shape VLPs from TomoNet and Relion tutorial. **e,** A zoom-in view of an irregularly shaped lattice. Coloring is based on surface curvatures at the point of each subunit.

### Application to focused ion beam (FIB)-milled cellular sample: the S-layer lattice of prokaryotic cell

We validated TomoNet’s particle picking capability by processing one tomogram of FIB-milled *Caulobacter crescentus* cells from EMD-23622^50^. The S-layer functions as a component of the cell wall covering the cell body. Thus, its lattice geometry is typically defined by the shape of cells (Figure 8a). The pleomorphic shape of *C. crescentus* cell in variable sizes, with the low contrast shown in this tomogram, hindered locating subunits on the S-layer lattice and raised difficulty for efficient particle picking on its S-layer lattice (Figure 8a).

**Figure 8.**
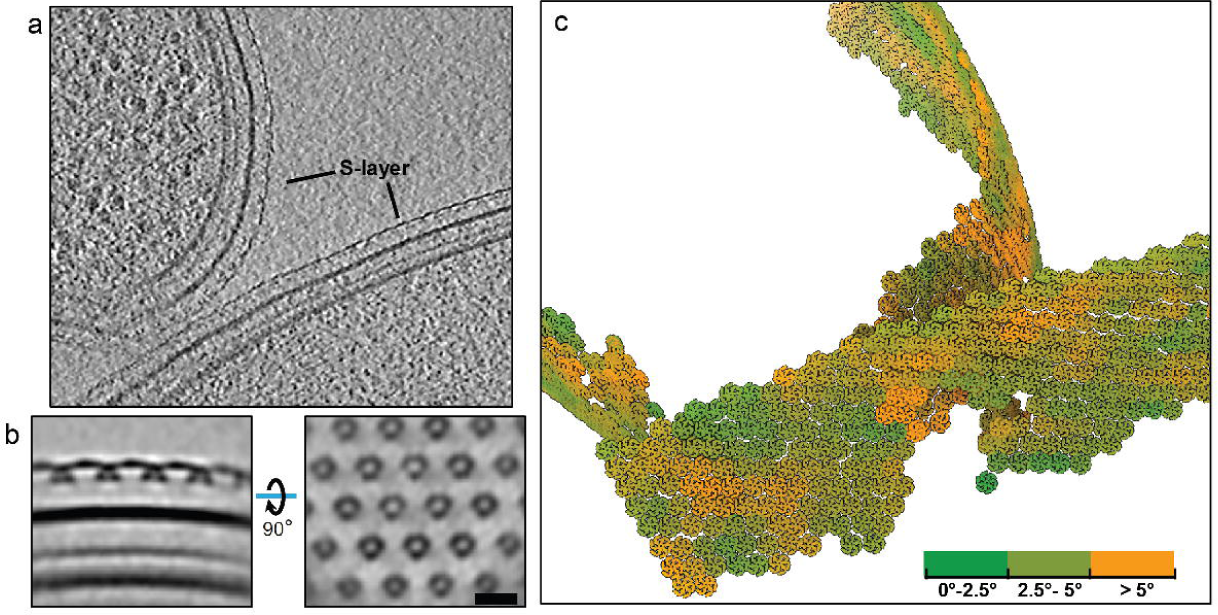
TomoNet application to S-layer structure in FIB-milled cellular sample. **a,** A tomographic slice view shows two *C. crescentus* cells in a FIB-milled lamella. **b,** Orthogonal slice views of the averaged density map generated in TomoNet, showing the hexagonal distribution of S-layer inner domains. Scale bar is 20 nm. **c,** Visualization of S-layer lattices generated by placing back hexamer subunit maps simulated from PDB: 6P5T. Coloring is based on surface curvatures at the center of each subunit.

TomoNet overcame the above challenges by utilizing the hexagonal configuration of S-layer lattices. With a minimal manual input, “Auto Expansion” picked over a thousand hexamer S-layer subunits. The intermediate STA result clearly reveals the hexagonal distribution of S-layer inner domains (Figure 8b). Visualization of S-layer lattices also shows that the picked particles were arranged in the expected hexagonal pattern, confirming the reliability and applicability of TomoNet as a particle picking tool (Figure 8c) and its broad application to structure determination of prokaryotic and archaeal cell walls^45,51^.

### Application to *in vitro* assembled arrays: nuclear egress complex (NEC) lattice

We further validated TomoNet as an integrated high-resolution STA pipeline and an efficient particle picking tool by processing samples containing NEC lattices within budded vehicles. Nuclear egress is a pivotal step in herpes virus replication, driven by NEC and responsible for translocating nascent viral particles from nucleus to cytoplasm. In our reported dataset^52^, NEC heterodimers budded into large vesicles with diameters ranging from 100 nm to 500 nm, forming beehive-like lattices on the inner surface of these vesicles (Figure 9a, b). Because of their large sizes, noticeable compressions were observed during the sample freezing, reshaping the vesicles and NEC lattices from spherical to flattened disk shapes (Figure 9a, b). This conformational change was a consequence of the limitation in ice thickness imposed by cryoET, which restricts the sample thickness to approximately 250 nm, consequently posing challenges for particle picking.

**Figure 9.**
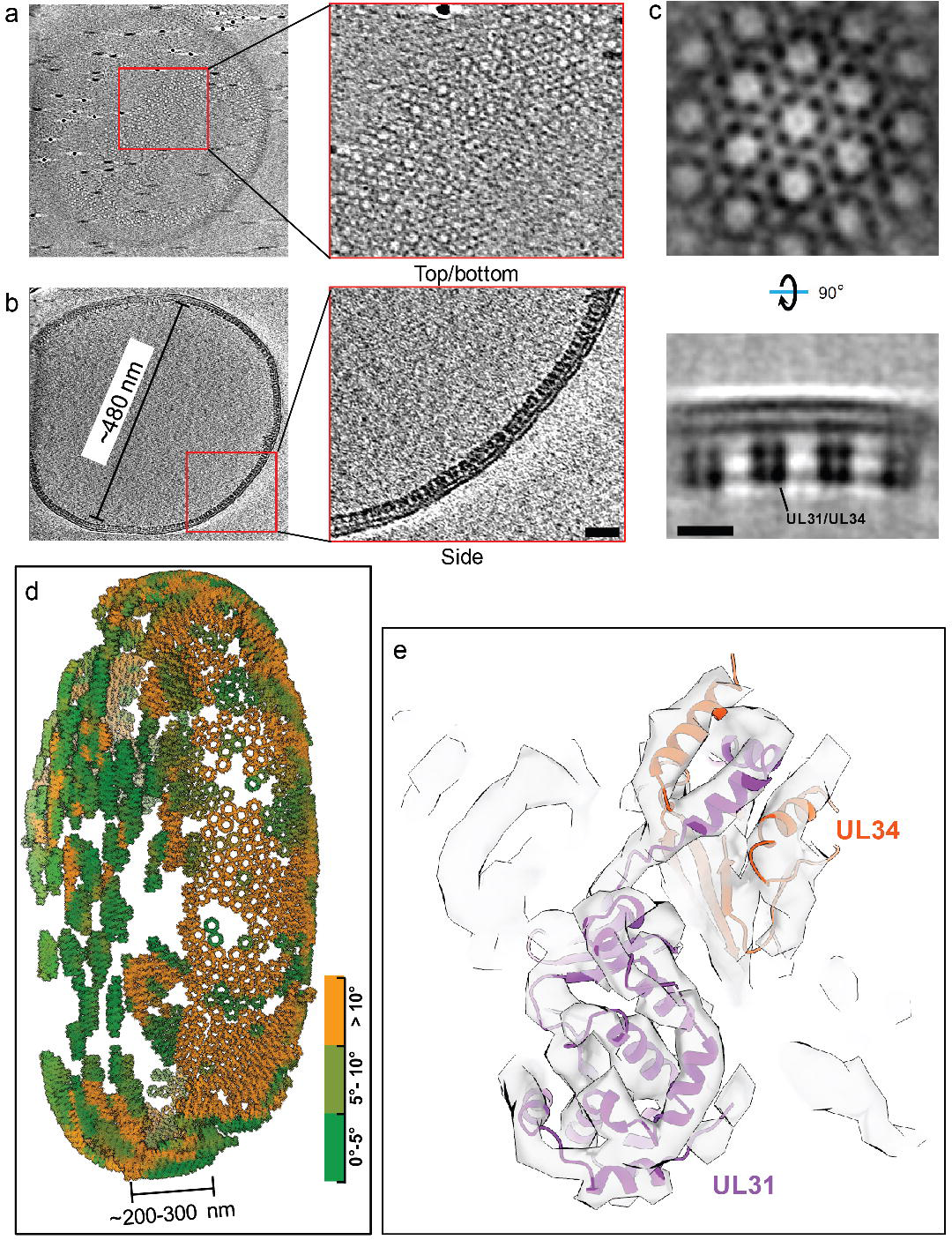
TomoNet application to *in vitro* assembled NEC-bound membrane. **a, b,** Tomographic slice views show a large NEC lattice; the insets show different views of NEC hexamer subunits. Scale bar is 20 nm. **c,** Orthogonal slice views of an averaged density map generated in TomoNet show that NEC hexamer subunits consist of UL31/UL34 heterodimers. Scale bar is 10 nm. **d,** Visualization of an NEC lattice generated by placing back averaged maps shows that the large vesicle is compressed into a disk-like shape. The compression caused by sample freezing stretched the lattice, making it flat and split at the air-water surface. Coloring is based on surface curvatures at the center of each subunit. **e,** Atomic model of the UL31/UL34 heterodimers fits into the final averaged map, with all helices well resolved.

TomoNet successfully picked NEC hexamer subunits following the topology of lattices. The intermediate STA result generated in TomoNet already showed the six heterodimers within one hexamer subunit (Figure 9c). With these picked particles, high-resolution 3D classifications and refinements were carried out to obtain a final reconstruction of NEC hexamer subunit at 5.4 Å resolution, without preferred orientation bias (Figure 9c, d and Figure 6c), and all the helices were well resolved (Figure 9e). Visualization of subtomograms placing back shows that the large vesicle was compressed during sample freezing which stretched the NEC lattice, making it appears flat and split at the air-water interface, while the middle part of the lattice appears to be more curved.

### Application to other types of arrays and free-floating particles

The above examples show how TomoNet’s ability to locate particles arrays arranged on flexible spheres (HIV), cell surfaces (S-layer) and nuclear membranes (NEC), which can be considered as topologically 2D lattices. In our published work of various cryoET structures, TomoNet has also been used to locate subtomograms arranged on flexible filaments (*i.e.*, 1D arrays) such as the flagella of *Trypanosoma brucei*^22,42^ and the amyloid-like sheath protein on β-hoops of the prototypical archaeon, *Methanospirillum hungatei*^53^. In the case of 3D lattices, TomoNet has been also used to obtain the paraflagellar rod structure of *T. brucei*^21^. Since TomoNet has integrated packages and is designed for the entire cryoET and STA data processing pipeline, it can also be used as a general-purpose package for subtomogram averaging towards high resolution when particles are free floating and without local order. In the latter case, TomoNet would have the same limitation recognized for all other cryoET software packages, that is, high resolution is currently only achieved for large complexes, such as ribosomes.

## DISCUSSION

In this paper, we report the implementation and application of TomoNet and demonstrate its efficacy in particle picking across three distinct datasets featuring particles with varying lattice configurations. TomoNet stands out as the first software to exhaustively trace lattices following its inherent topology. This unique approach ensures that the particle picking results faithfully reflect *in situ* or *in vitro* lattice shape, providing valuable insights into how these lattices are formed by their constituent subunits. For HIV VLPs, TomoNet application enabled us to directly visualize the VLPs lattices and their defects potentially caused by the absence of pentamer subunits. Similarly, for the NEC dataset, TomoNet facilitated a more direct observation of lattice conformation changes resulting from the sample freezing process. Since vesicles in this dataset were too large to be compressed from a sphere into a disk-like shape, the lattice regions near the air-water interface became stretched and subsequently divided into smaller fragments. Moreover, TomoNet demonstrated its exceptional performance, even when dealing with datasets characterized by extremely low contrast. For instance, in the cellular S-layer tomogram of a lamella, S-layer subunits were nearly imperceptible to human observations. Therefore, “Auto Expansion“ excelled in particle picking without requiring denoising or contrast-enhancement algorithms.

Additionally, “AI AutoPicking“, the deep learning-based module, demonstrated excellent performance on automatic particle picking, showing potential in handling a wide range of particle types even beyond those with lattice-like arrangements. Compared to the template matching-based “Auto Expansion”, “AI AutoPicking” has several advantages in particle picking. Firstly, it applies to particles situated on flexible lattices and those arranged in scattered patterns, such as cellular ribosomes. The neural network learns to pick by discerning 3D features of individual particles, and it does not require prior knowledge about lattice configuration. Secondly, it utilizes GPUs for fast convolution operations, enabling particle prediction in just several minutes for each tomogram. Thirdly, it does not require the “seed” particles used in “Auto Expansion”, which further reduces human efforts by approximately 5-15 minutes per tomogram. This is especially beneficial for processing extensive tomography datasets with hundreds of tomograms. However, comparing their final output particles, “AI AutoPicking” typically picks fewer particles than “Auto Expansion” because it misses certain particles on the flexible lattices. Thus, these two modules are complementary to each other and can be incorporated to further explore these missing particles.

Regarding the pipeline design, each module within TomoNet is designed to be highly independent, ensuring flexibility for integrating future methods and third-party packages. This adaptable framework positions TomoNet as a platform of choice for other developers to build their own innovations. At present, TomoNet is primarily tailored for integration with the Relion-related pipeline. However, it can accommodate specific demands and can be extended to integrate other pipelines, including emClarity^17^, EMAN2^4^, M^54^, and others in the future. In summary, TomoNet significantly simplifies the overall process for users in managing and monitoring every step of the complete cryoET and STA pipeline. Its user-friendly GUI design notably reduces the entry barrier for newcomers to the fast-emerging cryoET field. The particle picking modules of TomoNet provide a general solution for particles organized in lattice-like arrangements, ensuring both accuracy and efficiency, thereby facilitating the high-resolution STA pipeline.

## METHODS

TomoNet is an open-source software package developed using Python. It follows a highly modularized architecture with each module responsible for specific tasks in a typical cryoET and STA data processing pipeline. Modules in TomoNet mainly cover the upper stream of the cryoET and STA pipeline including procedures of motion correction, tilt series generation, tomogram reconstruction, CTF estimation and particle picking, while leave the high-resolution 3D refinement to established software package like Relion (Figure 1). The design of a modern GUI, established with PyQt5 platform, enhances user-friendliness, and helps with tracking the processing progress (Figure 2). With table views, users can obtain a comprehensive overview of the entire dataset, facilitating direct and intuitive management for each tomogram (Figure 2).

### Implementation of modules for motion correction, tomogram reconstruction and CTF estimation

Motion correction, tomogram reconstruction, and CTF estimation related functions are organized into individual modules in TomoNet, with the integration of corresponding external software packages including MotionCorr2^36^, IMOD^37^ or AreTomo^38^ and CTFFIND4^39^, respectively. Since their codes are not rewritten in TomoNet, users have to install each of them before using the corresponding modules.

The “Motion Correction” module is used to correct bean-induced sample motion. It requires an input folder path that contains all the dose fractionated frames, then user can specify their MotionCorr2 parameters in the GUI. After clicking the “RUN” button, TomoNet will perform motion correction for all the input images and save the results in a separated directory. This module also allows on-the-fly motion correction during data collection.

The “3D Reconstruction” module comprises two sub-functions: “TS Generation” and “Reconstruction”. Within “TS Generation”, users can readily assemble tilt series for each tomogram from the previously generated motion corrected images. It provides advanced options for data cleaning, such as setting a minimum acceptable number of tilt images for a tomogram, removing duplicate images at the same tilt angle by excluding images with older time stamps. The “Reconstruction” tab automatically reads and lists all tomograms in a table view, with essential information, such as tilt image number and alignment errors, and action buttons for restart, continue and delete individual tomogram reconstruction process. This simplifies the assessment of reconstruction results and facilitating tomogram reconstruction management.

The “CTF Estimation“ module is used for the tilt series defocus estimation, with support of parallel processing using multiple CPUs. Its outcomes are also listed in a table view with visualization features, such as displaying defocus at 0 degree and plotting the defocus distribution across all tilt angles.

### Implementation of the “Manual Picking” module

The “Manual Picking” module is designed for general management of manual particle picking, especially for the preparation of “seed” particles required in “Auto Expansion”. IMOD stalkInit picking criteria is implemented to define the Y-axis for each particle with 2 points, and the center in between them. In the example of HIV dataset, 5-10 particles were manually picked as the “seed” particles for each VLP lattice, which only takes several minutes per tomogram (Figure 5a).

### Design and implementation of the “Auto Expansion” module

“Auto Expansion” consists of three steps as shown in Figure 2. “Generate tomograms.star” is used to generate a STAR format file that maintain information of tomograms and their associated “seed” particles to be applied in “Auto Expansion”. “Generate Picking Parameter” is used to set up parameters required for particle set expansion through the described iterative process. The parameters include angular search ranges and steps, translational search ranges and steps, a “transition list” (explained later), box size used in particle alignment, distance between neighboring repeating subunits, reference and mask map, cross-correlation threshold, etc. The “transition list” is customized by users to describe the targeting lattice configuration, with each transition denoted by *[sx, sy, sz]*, where *sx, sy and sz* are translational shifts from the center of “seed” particle to one of its neighbors along X, Y and Z-axis, respectively. Thus, “Auto Expansion” can use it to guide the search of “candidate” particles. These user defined parameters will then be saved into a JSON format file. “Run Particle Expansion” takes the above STAR and JSON format files as inputs to perform the iterative particle set expansion.

During the “Auto Expansion” processing, three directories will be generated for each tomogram. They are “*TomoName*” as the working directory for carrying out the current iteration, “*TomoName_cache*” that stores intermediate results from finished iterations, and “*TomoName_final*” that stores the final particle picking results. The iteration number of “Auto Expansion” is typically greater than one. However, “Auto Expansion” allows for some special usage cases. For example, in the scenario when users need to modify the particle picking setting such as a different cross-correlation threshold, user can generate the new picking parameter file, then execute “Run Particle Expansion” by setting the iteration number as 0. This prompts the program to skip the “candidate” searching steps, but just gather all intermediate results saved in “*TomoName_cache*” directories, then generate a new “*TomoName_final*” result.

### Design and implementation of the “AI AutoPicking” module

The “AI AutoPicking” module comprise 3 main steps, “Prepare Training Dataset”, “Train Neural Network” and “Predict Particles coordinates”. It uses supervised machine learning that requires users to provide ground truth, *i.e.*, tomogram with the associated particle coordinates files, for the model training. In this study, the ground truth data were prepared by “Auto Expansion”.

In “Prepare Training Dataset”, extracted subtomograms are used as inputs to the network training model for two reasons. Firstly, the size of tomogram used for picking is typically around 1000×1000×1000 voxels which is not applicable to be loaded in the GPU memory, but the size of extracted subtomograms is under 100×100×100 voxels. Secondly, it helps with increasing the number of training data pairs to avoid over-fitting during the network training. For the model output, the particle coordinates information was embedded into 3D binary segmentation maps, where the voxels associated with particles were set to 1, otherwise set to 0 (Figure 4a).

In “Train Neural Network”, the above extracted subtomograms paired with their associated segmentation maps are used to train a neural network model to be a binary classifier that predict whether a voxel is near the center of a particle. The network architecture employed is derived from the one used in IsoNet^46^ as it is well-suited for capturing generalized features of 3D objects (Figure 4b). Since the learning task is voxel-wisely binary classification, cross entropy loss function is used instead of minimum squared error (MSE). Equipped with one RTX 3080Ti graphic card, the training process can be completed swiftly within 1-2 hours if using the default parameters.

In “Predict Particles coordinates”, users can apply the trained model on the entire tomography dataset for particle coordinate prediction (Figure 4c). For each tomogram, TomoNet generate a predicted segmentation map first, then its particle coordinates information can be retrieved from the segmentation map by utilizing the hierarchical clustering algorithm from *scipy* module in Python.

### Implementation of tools within the “Other Utilities” module

The “Other Utilities” module consists of two sub-functions: “Recenter | Rotate | Assemble to .star file“ and “3D Subtomogram Place Back“ as useful tools for post particle picking processing. The first one allows users to assemble and convert the particle picking results into a STAR format file following the Relion4 convention, reset particles center to its symmetric center, and align the rotation axis to Relion Z-axis. The second one takes a user-provided STAR format file that contains particles information as input, then generates a ChimeraX^55^ session file for 3D subtomograms placing back and a clean version of STAR format file with “bad” particles removed. This not only allows users to validate the accuracy of particle picking before importing into Relion, but also enables direct observation of the distribution and configuration of subunits after the high-resolution 3D refinements, providing overall *in situ* lattice observations (Figure 7).

### Processing tomograms of HIV VLP dataset

The HIV VLP dataset was downloaded from the Electron Microscopy Public Image Archive (EMPIAR) with the accession code EMPIAR-10164^43^. Four tilt series, TS_01, TS_43, TS_45 and TS_54, were used in this study. Downloaded micrographs were loaded into the TomoNet pipeline to perform tilt series assembly, CTF estimation, and tomogram reconstruction using the WBP algorithm.

Four-time binned tomograms with 5.4 Å pixel size were used for further particle picking. Firstly, tomograms TS_01 and TS_43 were used for “seed“ particles preparation on 3 selected VLPs per tomogram, and an initial reference map was generated by averaging them in PEET. Secondly, one run of “Auto Expansion” was applied on the above two tomograms to get more particles, such as to refine the reference. Thirdly, with an improved reference, a new run of “Auto Expansion” was applied on the selected 3 VLPs in both tomogram (Figure 5b), then the particle picking result was used for neural network training in “AI AutoPicking”. Fourthly, after the particle prediction on all four tomograms with a trained model, “AI AutoPicking” produced 4,860, 3,704, 4,550 and 2,101 particles for tomograms TS_01, TS_43, TS_45 and TS_54, as shown in Figure 5c. Lastly, the predicted particles were input as “seed” particles for the final run of “Auto Expansion”, resulting in 5,765, 4,043, 5,006, and 2,838 particles for tomograms TS_01, TS_43, TS_45 and TS_54, which were imported into Relion to perform high-resolution refinements.

Following the same procedure carried out in the Relion4 tutorial together with TomoNet 3D classification, the Gag hexamer structure was resolved at 3.2 Å resolution with 13,558 particles from four tomograms. Resolution was calculated in Relion and on 3DFSC Processing Server^56^. The global resolution reported is based on the “gold standard” refinement procedures and the 0.143 Fourier shell correlation (FSC) criterion (Figure 6).

### Processing one tomogram of *C. Crescentus* S-layer

The FIB-milled *C. crescentus* data of one reconstructed tomogram was downloaded from Electron Microscopy Data Bank (EMDB) with the accession code EMD-23622^50^. This tomogram was directly used for “seed” particles preparation on two of the cells. Around 30 “seed” particles were manually picked and averaged using PEET to generate an initial reference map. “Auto Expansion” was applied on the “seed” particles for 5 iterations to get more particles such as to refine the reference map. With the improved reference map, another run of “Auto Expansion” was applied to the same “seed” particles for 15 iterations to search all particles on the outer surface of the cells, and finally yielded ∼1,500 S-layer particles of hexamer subunits (Figure 8c).

### Processing tomograms of NEC budding *in vitro*

The cryoET grid preparation and data collection were previously described^52^. Motion correction, tomogram reconstruction and CTF estimation were performed using TomoNet. Around 50-150 “seed” particles were manually picked for each tomogram. “Auto Expansion” were applied on a total of 35 tomograms and yield the ∼48,000 particles before Relion refinements. Following one round of 3D auto-refine job under four-binned pixel size and several rounds of 3D auto-refine jobs under two-binned pixel size and one round of 3D auto-refine under unbinned pixel size, together with TomoNet 3D classifications, the NEC hexamer structure was resolved at 5.4 Å resolution with totally 35,039 particles.

### 3D visualization

IMOD^37^ was used to visualize the 2D tomographic and segmentation map slices. UCSF ChimeraX^55^ was used to visualize the STA results and the lattices generated by 3D subtomogram place back. The atomic models were fitted into the density map using the “fit in map” tool in ChimeraX.

## AVAILABILITY

TomoNet code is available on Github website at https://github.com/logicvay2010/TomoNet, with a user manual. For the HIV VLPs dataset, the raw data was downloaded from the Electron Microscopy Public Image Archive (EMPIAR) with accession code EMPIAR-10164^43^, the Gag atomic model was downloaded from the Protein Data Bank (PDB) with accession code 5L93^43^. For the *C. Crescentus* S-layer dataset, the reconstructed tomogram was downloaded from the Electron Microscopy Data Bank (EMDB) with accession code EMD-23622^50^, and the subunit model was generated using atomic model with PDB accession code 6P5T^57^. The STA result of NEC hexamer is from EMDB with accession code EMD-40224^52^. The other data that support the findings of this study are available from the corresponding authors upon reasonable request.

## ACKNOWLEDGEMENTS

We thank Elizabeth Draganova and Ekaterina Heldwein for the NEC dataset. This project is supported by grants from the US National Institutes of Health (GM071940 to Z.H.Z.) and the National Science Foundation (DMR-1548924 to Z.H.Z.).

## AUTHORSHIP CONTRIBUTIONS

HW and ZHZ initialized and ZHZ supervised research; HW wrote the code and developed the software GUI with help from SL; HW, SL and XY tested the software on different datasets; HW, and ZHZ wrote the manuscript; JZ and XY assisted the manuscript writing; all authors reviewed and approved the paper.

## Movie Legends

**Movie 1, A spherical VLP consisting of hexamer Gag subunits, colored by local surface curvature. “bad” particles with wrong alignment are shown as red.**

**Movie 2, A VLP lattice with irregular shape.**

